# Human cortical high-gamma power relates to movement speed and is disproportionately reduced after stroke

**DOI:** 10.1101/2023.11.07.565934

**Authors:** Benjamin Haverland, Lena S. Timmsen, Silke Wolf, Charlotte J. Stagg, Lukas Frontzkowski, Robert Oostenveld, Jan Feldheim, Focko L. Higgen, Christian Gerloff, Robert Schulz, Till R. Schneider, Bettina C. Schwab, Fanny Quandt

## Abstract

Motor cortical high-gamma oscillations (60 to 90 Hz) occur at the onset of movement and are spatially focused over the contralateral primary motor cortex. Although high-gamma oscillations are widely recognized for their significance in human motor control, their precise function on a cortical level remains elusive. Importantly, their relevance in human stroke pathophysiology is unknown. Understanding the neurophysiological processes of motor coding could be an important step in improving motor recovery after stroke. We recorded magnetoencephalography data during a thumb movement speed task in 14 chronic stroke survivors, 15 age-matched control participants and 29 healthy young participants. Motor cortical high-gamma oscillations showed a strong relation with *movement speed* as trials with higher *movement speed* were associated with greater high-gamma power. Stroke survivors showed reduced cortical high-gamma power, surpassing the effect attributable to decreased *movement speed* in these participants. Even though motor skill acquisition was evident in all groups, it was not linked to high-gamma power. Our study is, to our knowledge, the first to quantify high-gamma oscillations after stroke, revealing a reduction in movement-related high-gamma power. Moreover, we provide strong evidence for a pivotal role of motor cortical high-gamma oscillations in encoding *movement speed*.

## Introduction

Gamma oscillations (25 to 140 Hz) have been found in various areas of the human brain^1^ and have been linked to neural communication^2,3^ as well as to a wide range of cognitive processes.^4^ At the onset of motor actions, spatially focused movement-related high-gamma oscillations (∼ 60 to 90 Hz) occur over the contralateral motor cortex^5^, which have been shown to be prokinetic.^6–10^ High-gamma oscillations have been recorded in motor cortical areas from extracellular recordings (single- or multi-unit, local field potentials (LFPs))^11,12^, electrocorticographic (ECoG) recordings^7,13^, as well as non-invasive magnetoencephalography (MEG)^6,14^ and electroencephalography (EEG).^15^ In particular, invasive cortical recordings showed that high-gamma oscillations contain information that allows for decoding position, velocity and predominantly movement speed.^13,16,17^

Although high-gamma oscillations seem to play a crucial role in human motor control, they have so far never been quantified in stroke survivors. As over 50% of all stroke survivors suffer from persistent hand motor deficits^18^, understanding the neurophysiological processes of motor coding after stroke is key for the development of interventions to improve recovery and motor performance.^19^ Moreover, the study of human populations with impaired motor function may shed light on the general role of movement-related high-gamma oscillations. For example, if high-gamma oscillations purely reflect movement kinematics, they should be proportionally scaled down with impaired motor function. If high-gamma oscillations rather reflect the intent to move or the cognitive effort, they would be expected to be preserved or even enhanced with restricted motor function.

Here, we rigorously characterize the role of movement-related high-gamma oscillations in motor control and motor skill acquisition. We acquired a combined data set consisting of MEG data measured during a thumb movement task, anatomical and diffusion-weighted magnetic resonance imaging (MRI) as well as neurological tests from stroke survivors, healthy age-matched control participants and healthy young participants. We hypothesize that motor cortical high-gamma oscillations are tightly linked to *movement speed* and explore their role in motor control in the chronic phase after stroke.

## Materials and methods

### Participants and setting

Three different groups of individuals were recruited within the study: (i) 16 well-recovered stroke survivors, (ii) 18 healthy control participants age-matched to the stroke survivors and (iii) 30 healthy young participants (age 18 to 35 years). Stroke survivors, with a first-ever clinical stroke at least 6 months prior to inclusion, who experienced a deficit of the upper extremity for at least 24 hours and showed a structural lesion on a clinical MRI or computer tomographic image were included in the study. Psychotropic medication was an exclusion criterion in all three groups. The number of recruited stroke survivors was based on prior comparable observational studies.^20,21^ All participants were right-handed according to the Edinburgh Handedness Inventory. One healthy young participant and three age-matched older controls had to be excluded because they did not fulfil all inclusion criteria retrospectively (neurological conditions, handedness). Two stroke survivors were excluded because of extensive artifacts in the MEG data due to metal implants. This resulted in 14 stroke survivors, 15 age-matched controls, and 29 healthy young participants for the remainder of the study.

Multimodal imaging was obtained, consisting of MEG measurements during the execution of a motor task, as well as structural MR imaging. Standardized clinical testing was acquired in stroke and age-matched control participants by means of the Fugl-Meyer Assessment of the Upper Extremity (UEFM), Action Research Arm Test (ARAT), Nine Hole Peg Test (NHPT), Box and Block Test (BBT), whole hand grip force, key grip force, National Institutes of Health Stroke Scale (NIHSS), modified Rankin Scale (mRS) and Mini Mental State Examination (MMSE).

Participants gave written informed consent and the study was performed according to the Declaration of Helsinki. The study was approved by the local ethics committee of the Medical Association of Hamburg (2021-10410-BO-ff).

### Motor Task

Participants were seated comfortably in the MEG system in front of a screen with their performing arm fixated in a splint, next to a two-button response box (Current Designs, Philadelphia, PA) that was placed in a position where the thumb of the participant was able to easily reach both buttons. The task consisted of alternately pressing the two buttons for a total of four button presses in reaction to a ‘ready – steady – go’ cue using the thumb while keeping the arm as relaxed as possible (Fig. 1A).

**Figure 1.**
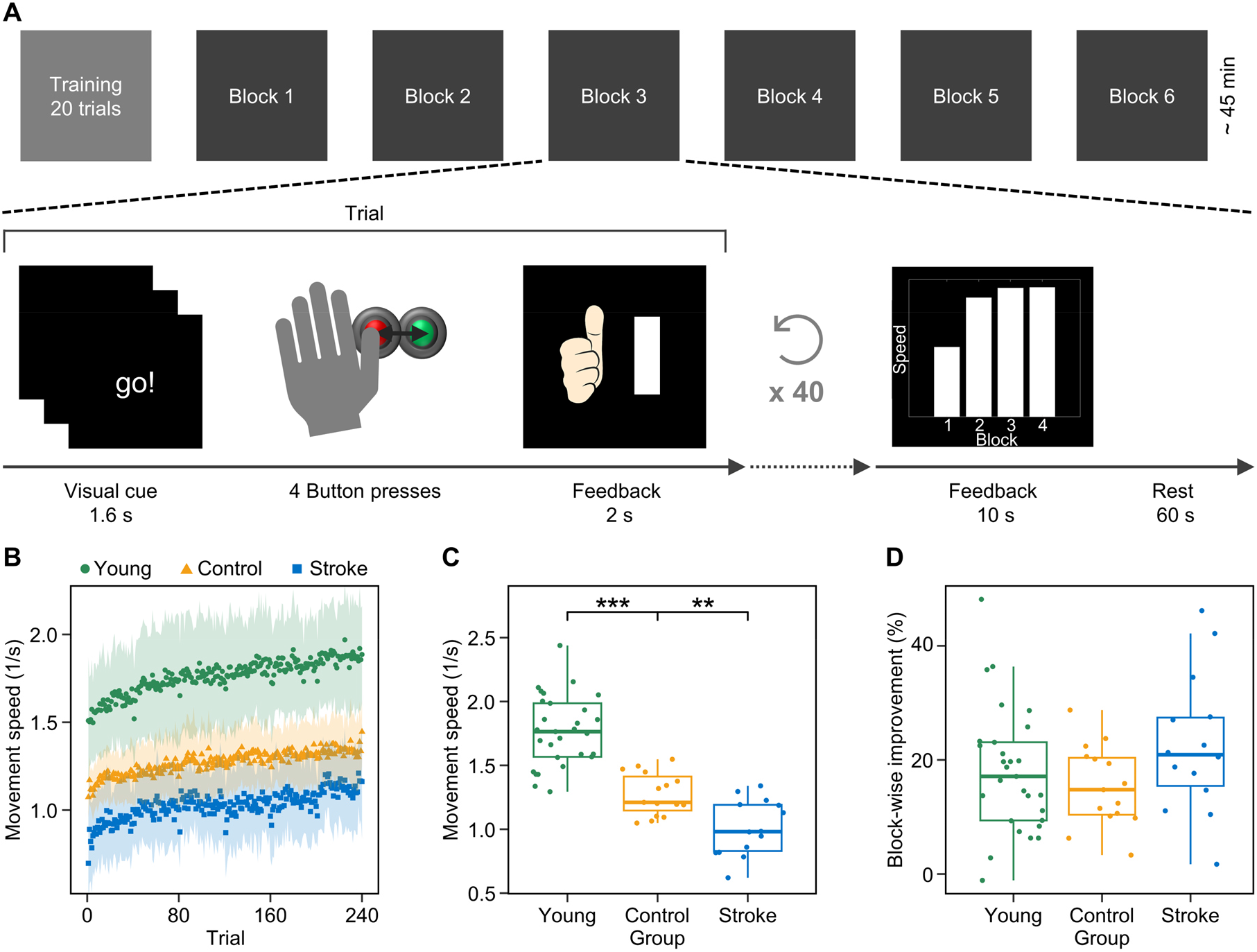
Motor task and behavioural data. (**A**) The motor task contained one training block with 20 trials and six measurement blocks with 40 trials each. In each trial, participants had to perform four alternating button presses as fast as possible in response to a visual cue. They received feedback on their *movement speed* in each trial. Additional feedback showing the block mean of *movement speed* was given after each block. There was a 1-minute rest period between blocks. (**B**) *Movement speed* for each trial averaged in each group. An increase in *movement speed* over the course of the trials was evident in all groups. Shaded areas indicate ± standard deviation. (**C**) Distribution of participant mean of *movement speed*. Asterisks indicate significant group differences. (**D**) Distribution of participant mean of *block-wise improvement. Block-wise improvement* represents the percentage *improvement* in *movement speed* from the first to the individual best block. There were no significant group differences in block-wise improvement. Significance markers: \*\**p* < 0.01, \*\*\**p* < 0.001

First, participants performed 20 trials to familiarize themselves with the task. Afterwards, they were instructed to perform the task as fast as possible using standardized instructions. Each participant performed six blocks with 40 trials each. After each trial, participants received feedback on their *movement speed* and the improvement relative to their mean *movement speed* in the current and the previous block. After each block, additional feedback was given in the form of a bar indicating the mean *movement speed* for each completed block. If participants pressed the button before the go-cue, took more than one second before initiating the first button press, or pressed an incorrect order of buttons, an error message was displayed.

There were differences in the experimental set-up between groups of young and control/stroke participants to account for behavioural differences and needs between these groups. Young participants performed the task with their left (non-dominant) hand, while stroke survivors used the hand contralateral to the lesioned hemisphere. The performing hand of age-matched control participants was chosen to match stroke survivors. Moreover, while in the young group lights in the measuring chamber were turned off, the light was kept on for stroke and older participants. Further, the size of the feedback bar (linear versus sigmoidal increase) was adapted in the age-matched control group and in stroke survivors to account for expected differences in *movement speed*. The task was run in MATLAB (R2021b, The MathWorks, Inc.) using Psychtoolbox-3.^22^

### Data acquisition

#### MEG and MRI data acquisition

MEG data were recorded with 275 axial gradiometers of a CTF-MEG system (CTF Systems, Coquitlam, Canada) with a sampling rate of 1200 Hz. The head position was continuously measured, and participants were given instructions to help them return to their initial head position after each block of the task.^23^ Electrooculogram (horizontal and vertical) and electrocardiogram were recorded via bipolar channels. Time points of button presses and visual stimuli were sent to the acquisition program via TTL triggers.

We acquired MRI data from 11 out of 14 stroke survivors, all 15 control participants and 21 out of 29 young participants. The remaining participants did not consent to MR-imaging. We acquired T1-weighted images in all groups and T2-weighted and multi-shell diffusion weighted images (DWI) in stroke and control participants using a 3 T Prisma MRI scanner (Siemens Healthineers, Erlangen, Germany) equipped with a 64-channel head coil. For anatomical imaging in stroke and control participants, a 3-dimensional magnetization-prepared rapid gradient echo sequence (repetition time (TR) = 2500 ms, echo time (TE) = 2.15 ms, flip angle 8°, 288 coronal slices with a voxel size of 0.8 × 0.8 × 0.8 mm^3^) was used. In young participants, the parameters of the sequence were slightly different (TR = 2300 ms, TE = 2.98 ms, flip angle 9°, 256 coronal slices with a voxel size of 1 × 1 × 1 mm^3^). T2-weighted images were acquired by using a fluid-attenuated inversion recovery sequence (TR = 9210 ms, TE = 92 ms, TI = 2500 ms, flip angle 140°, 70 axial slices with a voxel size of 0.9 × 0.9 × 2.0 mm^3^) for stroke lesion delineation. DWI were obtained covering the whole brain with gradients (b = 500, 1000 and 2000 s/mm^2^) applied along 96 non-collinear directions with the following sequence parameters: TR = 5000 ms, TE = 76 ms, slice thickness = 2 mm, in-plane resolution = 1 × 1 mm. For stroke survivor 14 the sequence parameters varied slightly: TR = 5000 ms, TE = 78 ms, slice thickness = 2 mm, in-plane resolution = 1 x 1 mm.

### Data analysis

Data analysis was conducted using the software package FieldTrip^24^ in MATLAB. Statistical analysis was performed in R version 4.1.3.^25^

#### Clinical and behavioural data

For UEFM, MMSE, ARAT, NHPT and BBT, we used the absolute test scores of the performing hand (UEFM: range 0 to 66, MMSE: range 0 to 30, ARAT: range 0 to 57, NHPT: pegs/second, BBT: blocks/min). Whole hand grip force and key grip force scores of the performing hand were divided by scores of the non-performing hand to obtain a ratio score. Within the motor task, *movement speed* was defined as the reciprocal of the time between the first and the fourth button press in each trial. *Improvement* was defined in two different ways:

i. on a trial level as the percentage increase in *movement speed* from the current to the next trial:

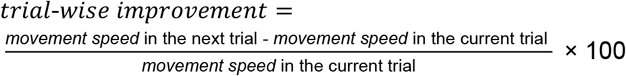
ii. across blocks as the change from first block to best block:

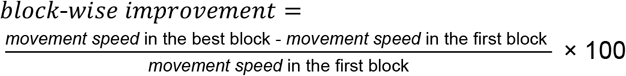

#### Processing of the MEG data

MEG data were bandpass filtered from 1 to 120 Hz and bandstop filtered at 49 to 51 Hz and 99 to 101 Hz (to eliminate the line noise artefact and its first harmonic) using a fourth-order Butterworth filter. Data were segmented into trials from -3.2 s before, up to +3.5 s after the time point of the first button press (0 s). Artifacts were carefully removed by a combination of visual artifact rejection as well as an independent component analysis, in which components representing eyeblinks, cardiac activity, or muscle artifacts were removed from the data. Error trials, trials in which participants took more than 3 s to complete all four button presses, as well as trials in which the movement duration (time from first to fourth button press) deviated more than three scaled absolute deviations from the individual block median were excluded, leaving on average 205.5 ± 16 (mean ± standard deviation) trials per participant in the young group (5960 trials in total), 191.5 ± 25.8 trials per participant in the stroke group (2681 trials in total) and 205.6 ± 14.6 trials per participant in the control group (3084 trials in total).

Source space activity was estimated using the dynamic imaging of coherent sources (DICS) beamforming method^26^ with regularization (λ = 5%). Head models, giving a geometrical description of the head, were computed based on individual T1-weighted MRIs (21 young participants, 11 stroke survivors, 15 control participants) or based on the Colin27 Average Brain^27^ (8 young participants, 3 stroke survivors) using the single-shell method.^28^ Source models, specifying the location of the sources in the brain volume, consisted of a regularly spaced 8 mm 3D grid warped on the individual or standard head model, respectively. At each location, a projection matrix describing signal propagation to the MEG sensors (the leadfield matrix) was computed. Time-frequency averaged cross-spectral density matrices for the movement period (0 to 0.6 s) and baseline period (−1.6 to -1 s) at 75 Hz with ± 15 Hz frequency smoothing (17 slepian tapers) were computed and source space activity was estimated at each grid point. To noise-normalize source power, the difference of power in the movement period and baseline period was divided by the baseline period for each trial in each participant. Brain regions were identified by masking of the source representation with the Brainnetome atlas.^29^ For each participant, the voxel with maximal power in the primary motor cortex (M1, regions A4hf, A4ul, A4ll, A4t, A4tl), premotor cortex (PMC, regions A6vl, A6cvl, A6dl, A6cdl) and supplementary motor area (SMA, region A6m) contralateral to their performing hand was detected. The power values of these voxels were used for further analyses.

To obtain a time-frequency representation of virtual channels, we computed real-valued DICS filters at 75 Hz ± 15 Hz frequency smoothing for the periods 0 to 1.5 s (task period) and -1.6 to -0.1 s (baseline period) and multiplied the filters of the voxels with maximal power in each region with the time-series data. In the resulting source level time-series data, we estimated power in the range 60 to 90 Hz (steps of 2 Hz and 0.05s, 0.25 s moving time window, 5 slepian tapers, resulting in ± 12 Hz frequency smoothing). We identified the high-gamma peak frequency for each participant by averaging the data within the time interval from 0 to 0.6 s over trials and determined the frequency with the highest power value. To identify the number of peaks in participant-level frequency-averaged time courses, we used the MATLAB function findpeaks to detect local maxima within a time interval starting at the first button press until 0.2 s after the last button press. Peaks were required to have a minimum width of 0.05 s and had to be separated by at least 21% of the movement duration.

#### Processing of the MRI data

##### Quantification of structural disconnection

Stroke lesions were delineated with ITK-SNAP^30^ and registered to a Montreal National Institute (MNI) 1 mm^3^ template. Hemispheres of all survivors with left-sided strokes were flipped to the other side. In one survivor, the stroke lesion was not detectable on the MRI anymore (Survivor 1). To calculate the structural disconnection for each survivor, lesion masks were analysed using the network modification NEMO2-toolbox.^31^ In brief, Change of Connectivity (ChaCo) values were calculated by overlaying the MNI-registered lesion masks onto a normative structural connectome derived from 420 healthy participants from the Human Connectome Project.^32^ The ChaCo scores resemble the number of disrupted streamlines divided by the total number of streamlines connecting a brain region to the rest of the network. We applied the Brainnetome atlas^29^ to compute ChaCo values for ipsilesional M1, ventral premotor cortex (PMv, regions A6vl, A6cvl), dorsal premotor cortex (PMd, A6dl, A6cdl) and SMA. The mean of ChaCo values of PMv and PMd was used as ChaCo of the PMC. In correspondence with pre-established studies, we removed all values below 0.02 due to noise artifacts.^31^

##### Fractional anisotropy of the corticospinal tract

Preprocessing and reconstruction of MRIs were performed using QSIPrep 0.16.1^33^, which is based on Nipype 1.8.5.^34^ Individual fractional anisotropy (FA) maps were computed on the preprocessed DWI imaged using MRTrix3^35^ function “dwi2tensor” and “tensor2metric”. The maps were registered on a FA template in MNI space (FMRIB58_FA standard space image https://fsl.fmrib.ox.ac.uk/fsl/fslwiki/FMRIB58_FA) using “antsregistration” (ANTs 2.4.0^36^). A CST template mask from the mesencephalon to the cerebral penduncle (MNI coordinates z = -25 to z = -20) was used to compute tract-related mean FA (for details refer to the original publication^37^). Mean CST-FA values of the hemisphere contralateral to the performing hand (affected hemisphere in stroke survivors) were divided by the mean CST-FA values of the hemisphere ipsilateral to the performing hand to obtain ratio scores. See Supplementary material for more details.

### Statistical analysis

#### Group differences

Differences in stroke survivor characteristics, *movement speed, block-wise improvement*, high-gamma power and high-gamma peak frequency were assessed using two-tailed *t*-tests and Wilcoxon rank-sum tests as appropriate.

#### High-gamma power compared to baseline

Two-tailed paired *t*-tests were calculated in each source voxel between movement period and baseline and corrected for multiple comparisons via a cluster-based permutation analysis^38,39^ with 1000 randomizations and a threshold for clustering of *α* = 0.01. Data from participants performing the task with the right hand were flipped across hemispheres prior to the analysis.

#### High-gamma power matched for movement speed

To evaluate whether group differences in high-gamma power are solely due to differences in *movement speed* or also due to an effect of group, we matched each trial of each stroke survivor to a trial with the same *movement speed* (± 1%) of a control participant performing the task with the same hand as well as each trial of each young participant to a trial of a control participant. Trials were then averaged within the participants and the group difference was assessed with a one-tailed *t*-tests with the alternative hypothesis being that stroke survivors have less high-gamma power than control participants and that control participants have less high-gamma power than young participants. Given the random selection of trials in the matching process, we performed the analysis 1000 times and used the median *p*-value as the indicator of significance.

#### Modelling the association of behavioural data, structural MRI and high-gamma oscillations

Linear mixed-effects models were calculated across groups for each motor region (M1, PMC) to assess the association between (i) high-gamma power and *movement speed* as well as (ii) high-gamma power and *trial-wise improvement*, controlled for *movement speed*. Additionally, in case of (i) high-gamma and *movement speed*, models were calculated for each group separately. Models included random slopes for *movement speed* and random intercepts for participant. In case of model singularity, the random slope was dropped from the model (*movement speed* model in the stroke survivor group in both motor regions, *trial-wise improvement* model across groups in the PMC). Further, linear regression models were fitted to (iii) evaluate the relation of high-gamma power and *movement speed* on participant-level, (iv) high-gamma peak frequency and *movement speed*, (v) high-gamma power and *block-wise improvement*, controlled for *movement speed* in block 1 and (vi) high-gamma power and clinical scores, ChaCo or CST-FA. Performing hand and group were added as a fixed effect in models with multiple levels of those variables. To ensure homoscedasticity and linearity of the residuals, high-gamma power was transformed to log(100+high-gamma power) in models (i) and (ii).

Furthermore, linear models were computed for activity at each voxel in source space to assess the topography of the relation between high-gamma power and *movement speed* or *block-wise improvement*. To circumvent the problem of multiple comparisons, we applied a cluster-based permutation analysis^38,39^ with 1000 randomizations of the assignment between high-gamma power and the variable of interest while keeping the assignment of group and performing hand to high-gamma power fixed. The threshold for clustering was set to *α* = 0.01. The summed *t*-values of the variable of interest within each cluster were compared to the distribution of summed *t*-values within clusters after randomization to compute the final *p*-value.

#### Number of high-gamma peaks

We modelled the dependent variable number of high-gamma peaks with independent variables group, *movement speed* and performing hand using cumulative link models (CLM, ordinal package) with a flexible logit link function for the regions M1 and PMC.

Visual inspection of residual plots did not reveal any obvious deviations from homoscedasticity or normality, proportional odds assumption held true in the CLMs. In linear mixed-effects models and CLMs, significance was obtained by likelihood ratio tests of the full model against a reduced model without the effect in question. In the case of multiple comparisons, *p*-values were corrected accordingly using the false discovery rate. Full statistical test and model results are provided in a tabular form in the Supplementary material.

## Results

### Participants

Fourteen stroke survivors (age = 65.4 ± 9.3 (mean ± standard deviation) years, 7 females, 8 with a lesion of the right hemisphere, performing hand: 8 x left hand, 6 x right hand), 15 age-matched control participants (age = 64.5 ± 8.4 years, 7 females, performing hand: 10 x left hand, 5 x right hand), and 29 young participants (age = 25.4 ± 4.6 years, 13 females) passed inclusion criteria. Stroke survivors and control participants showed no signs of cognitive impairment according to the MMSE (stroke: mean 28.9 points, range 27 to 30 points; control: mean 29.7 points, range 28 to 30 points).

### Clinical scores and structural imaging

Stroke survivors mostly showed only mild impairment of the upper extremity (Table 1, mean UEFM = 61.1), with a significant difference between stroke and control participants in UEFM (Fig. 2D), ARAT, NHPT and BBT (UEFM: *p* < 0.001, ARAT: *p* = 0.004, NHPT: *p* = 0.02, BBT: *p* = 0.001) but no significant difference in whole hand (*p* = 0.08) and key grip force ratio (*p* = 0.69). CST-FA values were not significantly different between stroke and control participants (Fig. 2C, *p* = 0.27). Stroke survivors had a mean ChaCo of 0.19 (range 0 to 0.59) in M1 and 0.14 (range 0 to 0.55) in PMC. Most frequently disrupted streamlines from M1 were the ones targeting ipsilateral motor areas (Fig. 2B).

**Table 1.** Clinical and demographic data of stroke survivors.

| ID | Sex | Age | Lesion side | TAS (months) | Stroke location | MRS | NIHSS | UEFM | ARAT | Grip force | KG force | NHPT | BBT |
| --- | --- | --- | --- | --- | --- | --- | --- | --- | --- | --- | --- | --- | --- |
| 1 | F | 70 | L | 52 | TC, CR | 1 | 0 | 65 | 57 | 1.11 | 1.16 | 0.39 | 50 |
| 2 | F | 79 | R | 8 | TC, CR, CI | 3 | 2 | 46 | 44 | 0.09 | 0.60 | 0.08 | 26 |
| 3 | M | 74 | R | 153 | BG, CI, CR | 2 | 1 | 50 | 54 | 0.51 | 0.60 | 0.20 | 33 |
| 4 | F | 71 | R | 52 | BG | 1 | 1 | 63 | 57 | 0.97 | 1.07 | 0.35 | 43 |
| 5 | F | 74 | R | 43 | PLCI | 1 | 0 | 60 | 56 | 0.89 | 0.80 | 0.38 | 47 |
| 6 | F | 43 | L | 38 | PLCI | 0 | 0 | 66 | 57 | 1.59 | 1.26 | 0.60 | 82 |
| 7 | F | 73 | L | 39 | MED | 1 | 1 | 58 | 56 | 0.92 | 0.88 | 0.31 | 49 |
| 8 | M | 65 | L | 9 | BG | 1 | 0 | 66 | 57 | 0.93 | 1.28 | 0.45 | 71 |
| 9 | M | 61 | R | 9 | PLCI | 1 | 0 | 66 | 56 | 0.89 | 1.12 | 0.32 | 39 |
| 10 | M | 58 | L | 6 | CR | 1 | 0 | 64 | 57 | 1.05 | 1.00 | 0.41 | 52 |
| 11 | M | 57 | L | 51 | MED | 1 | 2 | 63 | 53 | 0.98 | 1.14 | 0.38 | 55 |
| 12 | F | 67 | R | 129 | PO | 0 | 0 | 64 | 57 | 0.97 | 1.40 | 0.41 | 58 |
| 13 | M | 62 | R | 9 | CR | 1 | 1 | 60 | 54 | 0.68 | 0.90 | 0.23 | 35 |
| 14 | M | 61 | R | 16 | PRE | 1 | 0 | 64 | 54 | 0.69 | 0.75 | 0.33 | 48 |
| Mean stroke | F: 7 | 65.4 |  | 45.6 |  | 1.1 | 0.6 | 61.1*** | 54.9** | 0.88 | 1.00 | 0.35* | 49.1* |
| SD stroke |  | 9.3 |  | 45.3 |  | 0.7 | 0.8 | 6.1 | 3.5 | 0.33 | 0.25 | 0.12 | 14.7 |
| Mean control | F: 7 | 64.5 |  |  |  | 0 | 0 | 65.7 | 56.9 | 1.18 | 0.97 | 0.44 | 65.9 |
| SD control |  | 8.4 |  |  |  | 0 | 0 | 0.7 | 0.5 | 0.7 | 0.09 | 0.06 | 7.7 |

TAS: time after stroke, KG: key grip, SD: standard deviation, TC: thalamocapsular, CR: corona radiata, CI: capsula interna, BG: basal ganglia, PLCI: posterior limb of the capsula interna, MED: media infarct, PO: pons, PRE: precentral gyrus; \* $p < 0.05$ , \*\* $p < 0.01$ , \*\*\* $p < 0.001$ , statistical test of stroke against control group. Values of MRS, NIHSS, UEFM, ARAT, NHPT (pegs/second) and BBT (blocks/minute) are absolute test scores of the affected hand (performing hand in control participants). Values of grip force and key grip force are ratios between the affected and not affected hand in stroke survivors and performing and not performing hand in control participants.

**Figure 2.**
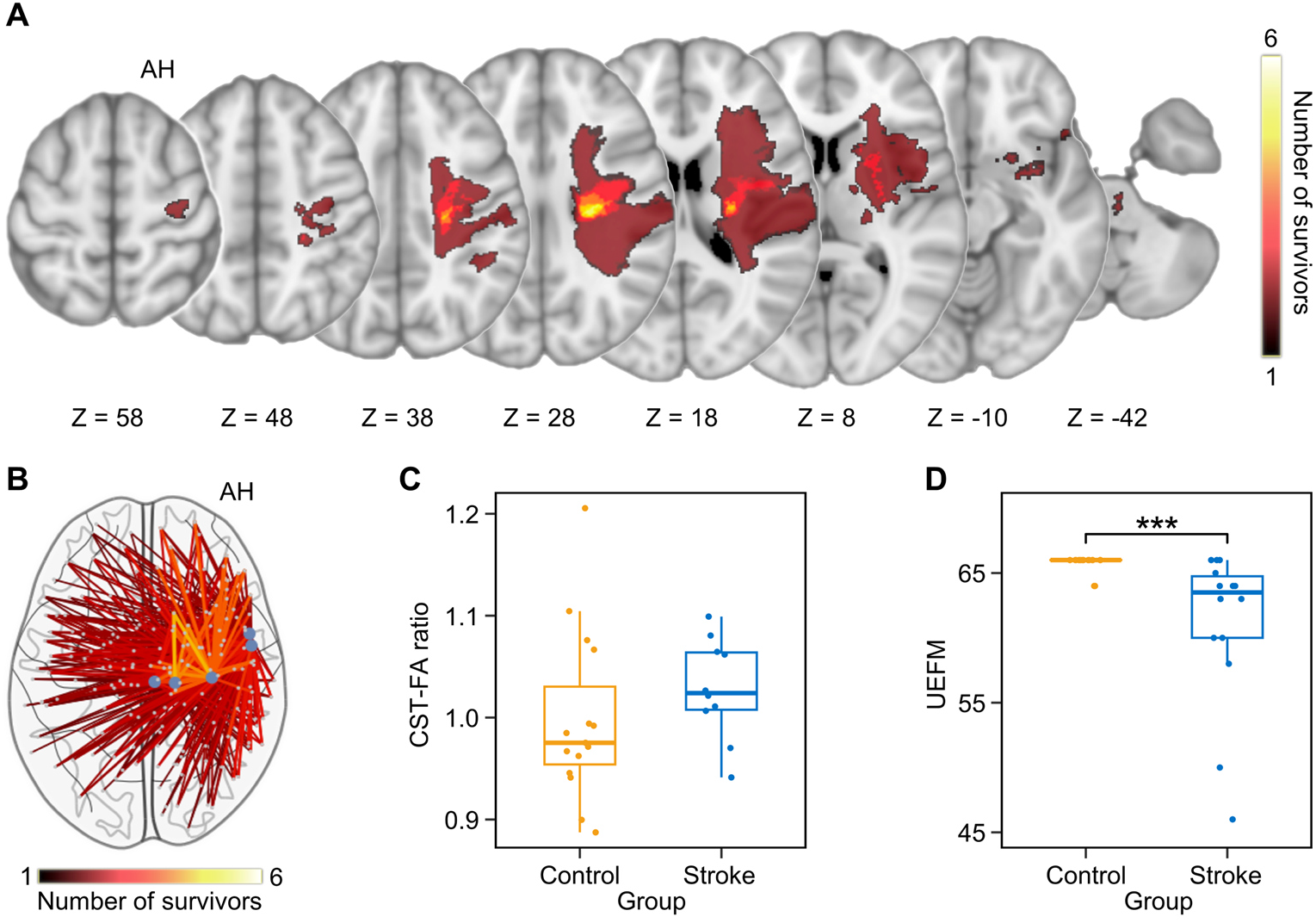
Stroke survivor characteristics. (**A**) Stroke lesions of the 10 survivors with available MRI lesion data, overlayed on a T1-weighted image in MNI standard space. Colour indicates the number of stroke survivors with a lesion at the voxel. Left-hemispheric lesions were flipped to the right hemisphere. (**B**) Structural disconnection of the 10 stroke survivors. Colour indicates number of stroke survivors with impaired connection between regions of M1 and the respective region. (**C**) Distribution of CST-FA ratios. (**D**) Distribution of UEFM-scores. Asterisks indicate the significant group difference. Significance marker: \*\*\**p* < 0.001

### Improvement in movement speed

On average, stroke survivors had a significantly lower *movement speed* than control participants (Fig. 1C, *p*_*cor*_ = 0.001; corrected for two comparisons), and control participants had a significantly lower *movement speed* than young participants (*p*_*cor*_ < 0.001; corrected for two comparisons). All three groups improved their average *movement speed* from block 1 to block 6 (Fig. 1B, stroke survivors: *p*_*cor*_ = 0.02, control participants: *p*_*cor*_ = 0.01, young participants: *p*_*cor*_ = 0.001; corrected for three comparisons). Comparing the initial and the individual best block, stroke survivors increased their average *movement speed* by 22.6 ± 12.2 % (mean ± standard deviation), control participants by 15.3 ± 7.1 % and young participants by 18.0 ± 10.9 %. *Block-wise improvement* showed no significant differences between the three groups, even without correction for multiple comparisons (Fig. 1D, stroke/control: *p* = 0.07, control/young: *p* = 0.33).

### Movement-related high-gamma power

We observed an increase in high-gamma power during the movement period (0 to 0.6 s, 60 to 90 Hz) relative to baseline with a maximum over the contralateral motor cortices (Fig. 3A). The increase of high-gamma power during movement was significant compared to baseline in each group (Supplementary Fig. 1, stroke survivors: *p*_*cor*_ = 0.006, control participants: *p*_*cor*_ = 0.003, young participants: *p*_*cor*_ = 0.003; corrected for three comparisons). The increase in high-gamma power occurred with movement onset and was most pronounced in M1 and PMC (Fig. 6C, Supplementary Fig. 2), prompting us to focus further analysis on these regions. Stroke survivors had significantly lower high-gamma power than control participants in the voxel with maximal power in regions M1 and PMC (Fig. 3B, M1: *p*_*cor*_ = 0.04, PMC: *p*_*cor*_ = 0.04; corrected for four comparisons) and control participants had significantly lower high-gamma power than young participants (M1: *p*_*cor*_ = 0.04, PMC: *p*_*cor*_ = 0.03; corrected for four comparisons). When matching performance (*movement speed*) and handedness in stroke survivors and age-matched controls (trials in each group 1206.1 ± 2.7 (mean ± standard deviation)), stroke survivors still had a significantly lower high-gamma power than control participants in both motor regions (Fig. 3C, M1: *p*_*cor*_ = 0.04, PMC: *p*_*cor*_ = 0.04; corrected for two comparisons). In contrast, the group difference in high-gamma power between young and control participants was not significantly different anymore after matching for *movement speed*, even without correction for multiple comparisons (trials in each group: 1470.8 ± 2.1, M1: *p* = 0.25, PMC: *p* = 0.15).

**Figure 3.**
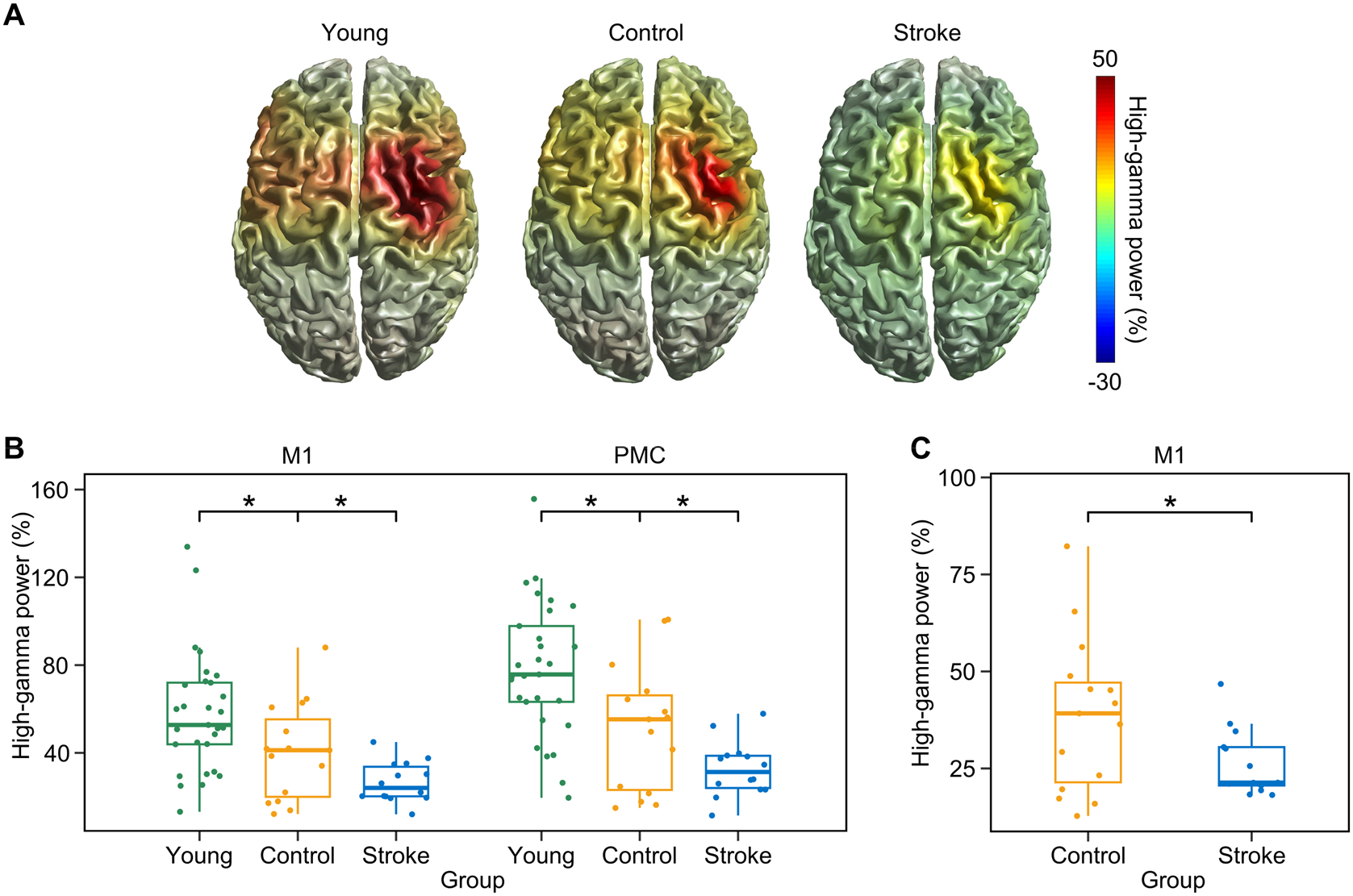
Spatial distribution and group differences of high-gamma power. (**A**) Group mean high-gamma power (0 to 0.6 s, 60 to 90 Hz) overlayed on a standard brain. Colour indicates the percentage increase in high-gamma power compared to baseline. Activity of participants that performed the task with the right hand was flipped across hemispheres. High-gamma activity is evident mainly in motor regions contralateral to movement. (**B**) Distribution of participant mean high-gamma power in M1 and PMC. Asterisks indicate significant group differences. (**C**) Group difference (control participants vs. stroke survivors) in high-gamma power when matched for *movement speed* and performing hand. Boxplots and dots show mean high-gamma power of each participant over 1000 repetitions of the matching process. Significance marker: \**p* < 0.05

### Relation of high-gamma power, structure and clinical scores

High-gamma power in stroke survivors from either motor region was not significantly related to the structural disconnection of M1 or PMC (ChaCo) (M1: *p* = 0.47, PMC: *p* = 0.79) or the structural integrity of the CST (CST-FA) (M1: *p* = 0.63, PMC: *p* = 0.45). Furthermore, there was no significant relation between high-gamma power in either motor region and residual motor function assessed by clinical scores (UEFM: M1: *p* = 0.74, PMC: *p* = 0.47; ARAT: M1: *p* = 0.09, PMC: *p* = 0.32; NHPT: M1: *p* = 0.87, PMC: *p* = 0.59; BBT: M1: *p* = 0.80, PMC: *p* = 0.56; grip force: M1: *p* = 0.81, PMC: *p* = 0.98; key grip force: M1: *p* = 0.82, PMC: *p* = 0.84).

### Relation of high-gamma power and movement speed

*Movement speed* significantly related to high-gamma power in linear mixed-effects models across all groups in both motor regions, as trials with higher *movement speed* were associated with greater high-gamma power (Fig. 4A, Supplementary Fig. 3A, M1: *p*_*cor*_ < 0.001, PMC: *p*_*cor*_ < 0.001; corrected for two comparisons). To investigate whether the effect of *movement speed* was driven by a single group, we modelled the relationship between high-gamma power and *movement speed* in each group individually. *Movement speed* was positively associated with high-gamma power in each group, however, after correction for multiple comparison only reached significance in the PMC of young participants (Fig. 4B, Supplementary Fig. 3C, stroke survivors: M1: *p*_*cor*_ = 0.14, PMC: *p*_*cor*_ = 0.14; control participants: M1: *p*_*cor*_ = 0.08, PMC: *p*_*cor*_ = 0.14; young participants: M1: *p*_*cor*_ = 0.07, PMC: *p*_*cor*_ = 0.03; corrected for six comparisons). Further evidence for the positive relation between high-gamma power and *movement speed* is provided by participant-level analyses across all groups. In those linear models, the same significant positive relation between *movement speed* and high-gamma power was evident (Fig. 4C, Supplementary Fig. 3B, M1: *p*_*cor*_ = 0.008, PMC: *p*_*cor*_ = 0.008; corrected for two comparisons). This participant-level linear model was stable enough to be computed in any region of the brain, even if the overall level of high-gamma power was low. Thus, to further infer on the spatial distribution of the association of high-gamma power and *movement speed*, we modelled the relationship for activity at each voxel. The positive relation was apparent even without preselection of the voxel with maximal power in motor regions. Importantly, the significant cluster only covered motor areas with a particular focus on the contralateral hand knob (Fig. 4D, *p* = 0.048).

**Figure 4.**
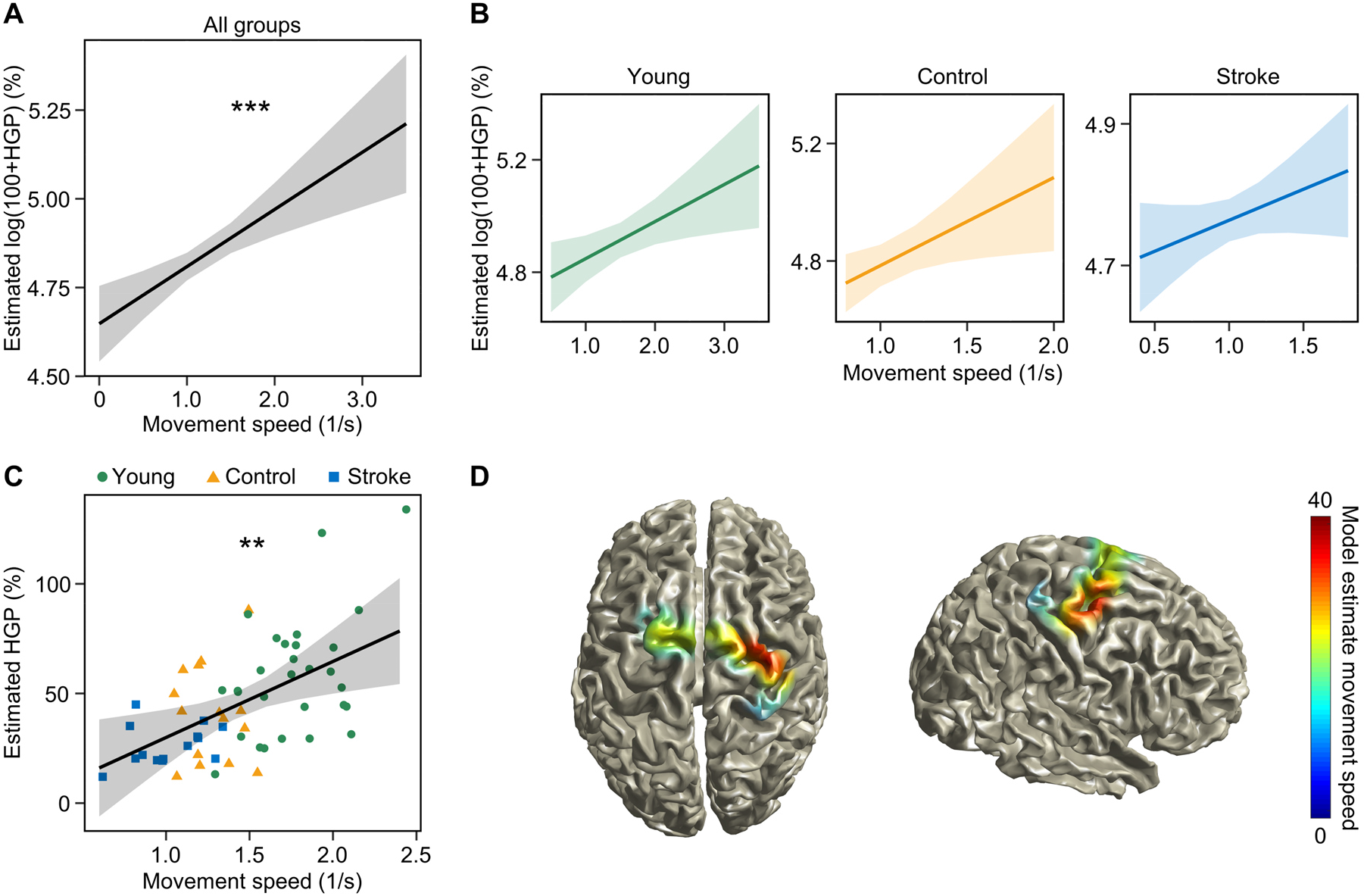
Relation between high-gamma power in M1 and movement speed. (**A**) Effect plot of the fixed effect *movement speed* in a trial-level linear mixed-effects model across groups for the relation between *movement speed* and high-gamma power. (**B**) Effect plots of the fixed effect *movement speed* in trial-level mixed-effects models for the relation between *movement speed* and high-gamma power. For each group individually, the direction of the relation is positive, although not significant. (**C**) Effect plot of the fixed effect movement speed in a participant-level linear regression model across groups for the relation between *movement speed* and high-gamma power. (**D**) Resulting cluster of a voxel-wise analysis for the relation of *movement speed* and high-gamma power. For activity in each voxel in the brain, a linear model between *movement speed* and high-gamma power was computed and results were tested for significance with a cluster-based permutation analysis. Colour indicates the model estimates of *movement speed*. Asterisks indicate significance of the effect *movement speed*. HGP: high-gamma power. Significance markers: \**p* < 0.05, \*\**p* < 0.01, \*\*\**p* < 0.001

### High-gamma peak frequency

High-gamma peak frequency significantly differed between young (M1: 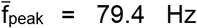) and control participants (M1: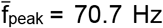), with significantly lower high-gamma peak frequency in the older controls (Fig. 5A, M1: *p*_*cor*_ < 0.001, PMC: *p*_*cor*_ < 0.001; corrected for four comparisons). High-gamma peak frequency was not significantly different between stroke survivors (M1: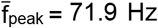) and control participants (M1: *p*_*cor*_ = 0.95, PMC: *p*_*cor*_ = 0.95; corrected for four comparisons). Linear regression models across groups showed no significant relation between *movement speed* and high-gamma peak frequency even without correction for multiple comparisons (M1: *p* = 0.15, PMC: *p* = 0.07).

**Figure 5.**
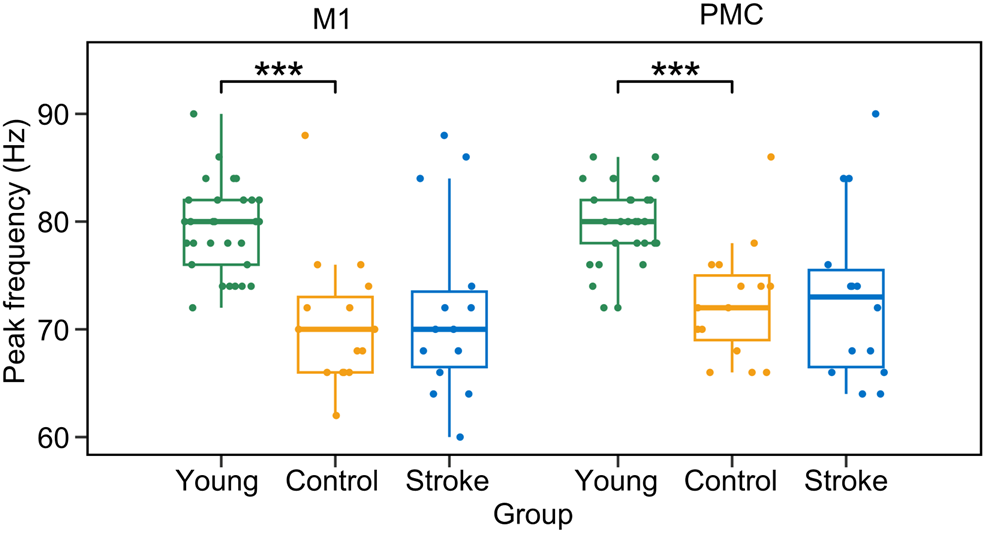
Characteristics of high-gamma peak frequency. Distribution of participant’s mean high-gamma peak frequency. Asterisks indicate significant group differences. Significance marker: \*\*\**p* < 0.001

### Number of high-gamma peaks

High-gamma power time courses revealed differences of high-gamma power evolution over time between the three groups (Fig. 6A). Young participants exhibited fewer high-gamma peaks over the time course of movement compared to control participants, the difference between control participants and stroke survivors was not significant (Fig. 6B, Supplementary Fig. 4A, control/young: M1: *p*_*cor*_ = 0.02, PMC: *p*_*cor*_ < 0.001; stroke/control: M1: *p*_*cor*_ = 0.40, PMC: *p*_*cor*_ = 0.58; corrected for four comparisons). The difference in number of high-gamma peaks across groups, however, turned out to be a result of differences in *movement speed*. When modelling the number of high-gamma peaks as a function of group and *movement speed*, we observed a significant effect of *movement speed* (Fig. 6D, Supplementary Fig. 4B, M1: *p*_*cor*_ < 0.001, PMC: *p*_*cor*_ = 0.006; corrected for two comparisons), but not group (M1: *p*_*cor*_ = 0.98, PMC: *p*_*cor*_ = 0.58; corrected for two comparisons). Hence, participants with faster *movement speed* had a significantly higher probability of having fewer high-gamma peaks. Specifically, in M1, if *movement speed* was greater than 1.88 /s (corresponding to ∼ 7.5 presses/s), the probability was highest that the participants would exhibit only one high-gamma peak. In contrast, if *movement speed* was below 0.62 /s (corresponding to ∼ 2.5 presses/s), the probability was highest that the participant would exhibit one individual high-gamma peak for each button press.

**Figure 6.**
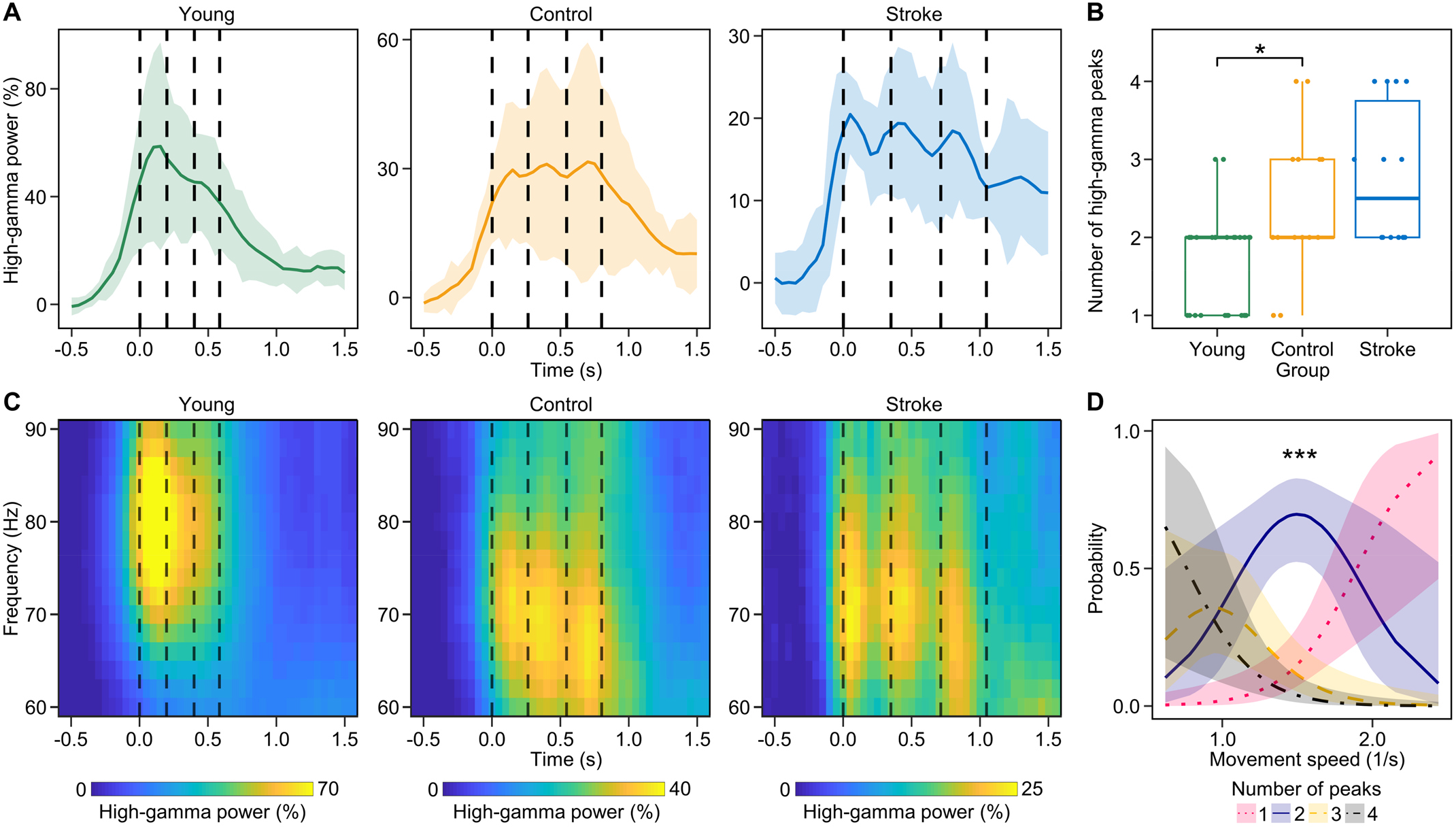
High-gamma power time course. Data shown are from the voxel with maximal power in M1, results are similar in PMC (Supplementary Fig. 4). (**A**) Group-level high-gamma power time courses. Dotted lines indicate average time points of button presses. Shaded areas indicate median absolute deviation. (**B**) Distribution of number of individual high-gamma peaks. Number of peaks was counted in participant mean of high-gamma power time courses. Asterisk indicates the significant group difference. (**C**) Group-level time-frequency spectrograms. The time point of the first button press corresponds to 0 s. Dotted lines indicate average time points of button presses. Note the different scaling of the colour bar in the different groups. (**D**) Effect plot of the independent variable *movement speed* in a cumulative link model that models the relationship between number of high-gamma peaks, *movement speed* and group with participant-level data from all three groups combined. The effect of *movement speed* was significant compared to a reduced model, the effect of group (not shown) was not significant. Significance markers: \**p* < 0.05, \*\*\**p* < 0.001

### Relation of high-gamma power and improvement

Neither *improvement* across trials, defined as the percentage increase in *movement speed* from the current to the next trial (*trial-wise improvement*) nor *improvement* across blocks, defined as the change from first block to best block (*block-wise improvement*) showed a significant association with high-gamma power (*trial-wise*: M1: *p* = 0.77, PMC: *p* = 0.48; *block-wise*: M1: *p* = 0.13, PMC: *p* = 0.38). In line, a cluster-based permutation analysis, modelling the association of activity in each voxel separately, showed no significant relation between high-gamma power and *block-wise improvement*.

## Discussion

Our study is the first to our knowledge to characterize cortical high-gamma oscillations with differing performance levels across age groups and in stroke survivors. We demonstrated that faster *movement speed* is robustly related to increased cortical high-gamma power over motor areas during movement onset. In contrast, *improvement* in performance did not significantly relate to high-gamma power. Moreover, with lower *movement speeds*, we observed distinct high-gamma peaks for each movement, while in faster movements, high-gamma activation became smeared over time. Importantly, even though *movement speed* showed the strongest association with high-gamma power, stroke pathology resulted in an additional decrease in cortical motor high-gamma power.

### Decreased cortical high-gamma power in stroke survivors

In humans, studies on movement-related high-gamma oscillations are restricted. Most reports on high-gamma oscillations have utilized invasive ECoG recordings, which are typically not available in stroke survivors. Scalp EEG, which can be measured easily in stroke survivors, can smear the focal high-gamma oscillations due to spatial blurring at the skull. MEG recordings, however, offer the opportunity to measure motor cortical high-gamma oscillations non-invasively and with high spatial precision in survivors with stroke pathology. In line with the literature in healthy cohorts, we observed high-gamma oscillations in chronic stroke survivors at the onset of contralateral movements, located over primary motor cortex and premotor cortices. We observed that chronic stroke survivors, even though exhibiting largely very mild hand motor impairment, showed less cortical high-gamma power compared to age-matched controls. This difference even remained significant after matching for *movement speed*. These results are paralleled by an existing study in rodents, in which the peri-infarct cortex of mice after stroke showed a deficit in low-gamma power under anesthesia.^40^ The disproportional decrease of high-gamma power in stroke survivors suggests that high-gamma power may not purely reflect movement kinematics. This suggestion is further supported by findings of significant high-gamma power in motor imagery.^41,42^ The decrease in high-gamma power was not related to clinical motor function scores or structural integrity of the corticospinal tract. Moreover, the structural disconnection of the primary motor cortex to other brain areas was not significantly related to the decrease of high-gamma power. The lack of a significant relationship between high-gamma power and structural measures in stroke survivors might possibly be due to the low number of survivors on whom we were able to acquire MRIs rather than the actual lack of a relationship, as corticospinal connectivity is related to motor impairment after stroke.^43^

### Topological organization of high-gamma oscillations

High-gamma oscillations are more spatially focused compared to lower frequencies^7^ and better aligned with somatotopy.^5,14^ In comparison, modulation of beta activity (13 to 30 Hz) may reflect a more general, widespread activation signal during and after movement with only limited specificity for the limb component that was moved.^9^ In simple motor tasks, high-gamma oscillations have been described to be localized to M1.^5–7^ With complex tasks, a more widespread cortical area is recruited^44^, consisting of PMd and PMv as part of the network for hand motor control.^45,46^ PMC is connected with both the arm area of the primary motor cortex and the spinal cord^47^ and is thought to be involved in connecting external cues with forthcoming movement.^45^ A reason for the observed elevated high-gamma power levels in PMC might be that our task required to constantly maximize motor performance. It may have thus involved a more complex network, including PMC, compared to simpler tasks like finger abductions or fist clenching.

### The role of high-gamma oscillations in motor control

High-gamma oscillations are thought to be prokinetic in nature and have been shown to strongly synchronize during active but not passive movements.^8,48,49^ At the onset of movement, in addition to the motor cortex, also basal ganglia and thalamus exhibit gamma band oscillations.^50–53^ Invasive recordings from the basal ganglia in patients with deep brain stimulation^41,48,54–56^ revealed that movement-related high-gamma power from the contralateral subthalamic nucleus and globus pallidus interna increased with movement amplitude, movement velocity and force production, stressing its potential significance in movement coding. Also, links to behaviour have been established in movement disorders, in which less high-gamma power in the contralateral subthalamic nucleus is associated with higher symptom severity in Parkinson’s disease (PD).^56^ Similarly, PD patients in the dopamine-depleted state show pathologically impaired movements and less high-gamma power.^54,55^ In line, dopaminergic medication, as a prokinetic drug, leads to increased high-gamma power in PD patients.^57^ Furthermore, patients suffering from hyperkinetic movements, such as medication-induced dyskinesias, present with pronounced high-gamma power.^58^ Recent studies have examined the subcortical-cortical communication in the gamma band and found that fast reaction times were preceded by enhanced subcortical spike to cortical gamma phase coupling^59^ underlining the role of gamma activity within the subcortical-cortical network for motor performance. In cortical invasive recordings, high-gamma oscillations up to very high frequencies (600 to 1000 Hz) contained decoding information on position, velocity and predominantly movement speed.^13^ Whereas single-unit activity in M1 is thought to mainly tune direction, larger neuronal populations as reflected by LFPs seem to represent movement speed.^16,17,60^ Here, for the first time using non-invasive recording in humans, we demonstrate a clear association of cortical high-gamma power (60 to 90 Hz) and *movement speed* across three groups with differing levels of *movement speed*.

### Time course of high-gamma oscillations

High-gamma oscillations have been shown to be most pronounced around movement onset, subsiding over the course of movement even when force production is maintained.^8^ Hence, high-gamma oscillations seem to encode, at least to some extent, kinematic variables or cognitive processes rather than purely active contraction of muscles. With repetitive movements, a distinct burst of high-gamma power has been described, with a tendency for individual gamma bursts to blur at a movement frequency of 3 Hz.^8^ We showed that the observed inter-subject difference in high-gamma peaks was driven by differences in *movement speed*, where with faster movements individual peaks blurred and the probability to exhibit only one high-gamma peak increased. This contrasts with observations for power at lower frequencies: beta-band power modulations have still been observed at a movement frequency of 4 Hz.^61^ Hence, for faster repeating movements, high-gamma oscillations may code the overall motor action rather than every single movement separately.

### Effects of high-gamma peak frequency

The range of motor high-gamma oscillations has been reported to be variable across studies, ranging from 75 to 100 Hz^7^ over 60 to 90 Hz^8^ to 55 to 375 Hz.^54^ In the hippocampal CA1 region, studies have described a slow-, mid- and fast-frequency gamma band. It remains unclear whether distinct high-gamma bands encode separate forms of motor processing. Over the lifespan, the high-gamma frequency range has been reported to vary depending on age with an increase from the very young child to adulthood.^62,63^ Here, we observe a decrease of the high-gamma peak frequency with age with significantly lower peak frequency in the older controls (79.4 Hz versus 70.7 Hz). These findings are in contrast to a previous study, which found no association between high-gamma peak frequency and age in a large cohort with a simple motor task.^64^ The reason for the differences might be the different analysis methods or discrepancies in motor tasks. Intra-participant gamma peak frequency has been reported to be consistent over recording sessions, over hemispheres and for movement of the same body part with differing movement vigour.^48,56,65^ Also in our study, high-gamma peak frequency did not significantly change with varying *movement speed*.

### The role of high-gamma power in motor skill acquisition

Recent work suggested that motor-related high-gamma oscillations may also be relevant for motor learning and plasticity^66–68^, and that motor skill acquisition is dependent on changes in local circuitry within M1.^69,70^ Some studies have shown an increase in motor performance (faster reaction times, learning of movement parameters) after high-gamma transcranial alternating current stimulation (tACS).^71–76^ Further, a study analysing transcranial magnetic stimulation-induced intracortical inhibition reported a decrease in intracortical inhibition after high-gamma tACS, correlating with improved motor learning scores.^77^ In our study, the level of motor skill acquisition was similar in stroke survivors compared to young and age-matched control participants, consistent with the notion that there is no significant difference in implicit motor learning in the affected hand after stroke compared to healthy controls.^78^ Hence, there seems to be potential for further rehabilitation to improve motor skills even in a cohort of mildly impaired stroke survivors. We investigated the relationship between high-gamma power and *improvement* in *movement speed* over time and did not find a significant association. Our findings suggest that high-gamma power does at least not strongly relate to motor skill acquisition in this dexterous finger movement task.

### Limitations

Several limitations should be considered for this study. First, the sample size of stroke survivors was limited. It is therefore possible that we missed findings because of restricted statistical power. Second, most stroke survivors exhibited only mild impairment, limiting generalizability to more severely impaired stroke survivors. Nevertheless, we were able to show a highly robust relationship between movement speed and high-gamma power and significant group differences between stroke and control participants. Third, as we specifically investigated thumb movements, the generalisability to movement of other parts of the upper extremity is unknown. Still, high-gamma oscillations are elicited by movements of different body parts in a similar manner^5,7,8^, which suggests that our findings may potentially generalize to other movement paradigms. Finally, we found no association between high-gamma power and motor skill acquisition, while it is important to note that our study misses any evaluation of long-term effects neglecting the learning-performance distinction.^79^ We therefore are only able to draw conclusions on short-term motor skill acquisition.

## Conclusion

In conclusion, characterizing cortical movement-related high-gamma oscillations, we found a strong positive relationship between *movement speed* and high-gamma oscillations within the motor network. For the first time, we quantified high-gamma oscillations in stroke survivors with motor impairment and observed a decrease in cortical motor high-gamma power going beyond the association to *movement speed*. We suggest that enhancing cortical high-gamma activity through neuromodulation could potentially serve as a therapeutic approach to improve motor performance in stroke survivors.

## Acknowledgements

We thank Karin Reimann and Christiane Reißmann for assistance in data recording and participant recruitment and Sophie Grigutsch for valuable discussions.

## Supplementary material

Supplementary material is available at bioRγiv online.

## Data availability

The data that support the findings of this study are available from the corresponding author, upon reasonable request.

