## Supplementary Material for "Human cortical high-gamma power relates to movement speed and is disproportionately reduced after stroke"

Benjamin Haverland[1,2], Lena S. Timmsen[1,2], Silke Wolf[1], Charlotte J. Stagg[3,4], Lukas Frontzkowski[1], Robert Oostenveld[5,6], Jan Feldheim[1], Focko L. Higgen[1], Christian Gerloff[1], Robert Schulz[1], Till R. Schneider[2], Bettina C. Schwab[7,2,#,*], Fanny Quandt[1,#,*]

1 Department of Neurology, University Medical Center Hamburg-Eppendorf, 20246 Hamburg, Germany

2 Department of Neurophysiology and Pathophysiology, University Medical Center Hamburg-Eppendorf, 20246 Hamburg, Germany

3 Wellcome Centre for Integrative Neuroimaging, FMRIB, Nuffield Department of Clinical Neurosciences, University of Oxford, OX3 9DU Oxford, UK

4 Research Council Brain Network Dynamics Unit, Nuffield Department of Clinical Neurosciences, University of Oxford, OX3 9DU Oxford, UK

5 Radboud University, Donders Institute for Brain, Cognition and Behaviour, 6525EN Nijmegen, The Netherlands

6 NatMEG, Karolinska Institutet, 171 77 Stockholm, Sweden

7 Biomedical Signals and Systems, Technical Medical Centre, University of Twente, 7522NB Enschede, The Netherlands

### These authors contributed equally to this work

* Corresponding authors: Fanny Quandt, University Medical Center Hamburg-Eppendorf, Martinistraße 52, 20246 Hamburg, Germany,; Bettina C. Schwab, University of Twente, Drienerlolaan 5, 7522NB Enschede, The Netherlands,

#### Supplementary Figures

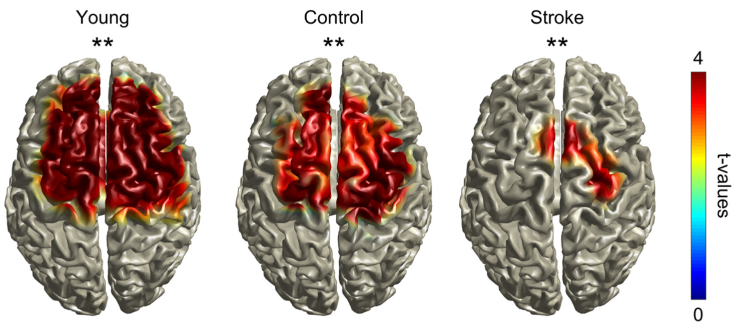


**Supplementary Figure 1 Difference in high-gamma power during movement compared to baseline.** Resulting clusters of a cluster-based permutation analysis for the difference in high-gamma power (60 to 90 Hz) in the movement period (0 to 0.6 s) compared to baseline. For activity in each voxel in the brain, a paired t-test between the movement period and baseline was calculated and results were tested for significance with a cluster-based permutation analysis. Colour indicates *t*-values. Asterisks indicate significance. Significance marker: **$p < 0.01$

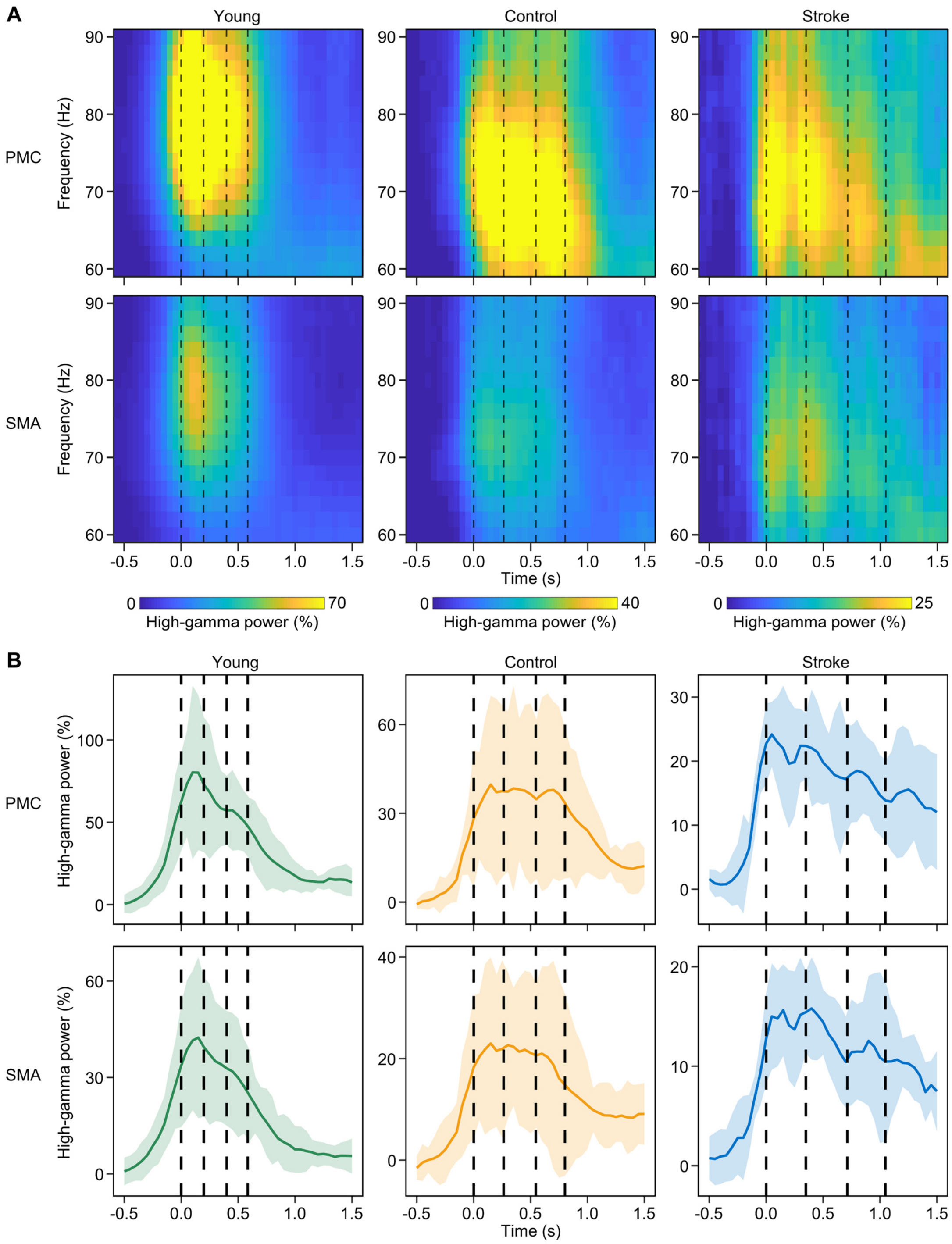


**Supplementary Figure 2 High-gamma power in PMC and SMA.** (**A**) Group-level time-frequency spectrograms. The timepoint of the first button press corresponds to 0 s. Dotted lines indicate average time points of button presses. Note the different scaling of the colour bars in the different groups. High-gamma power in M1 (Fig. 6) and PMC was higher than in SMA. We therefore excluded SMA from further analysis. (**B**) Group-level high-gamma power time courses in PMC and SMA.

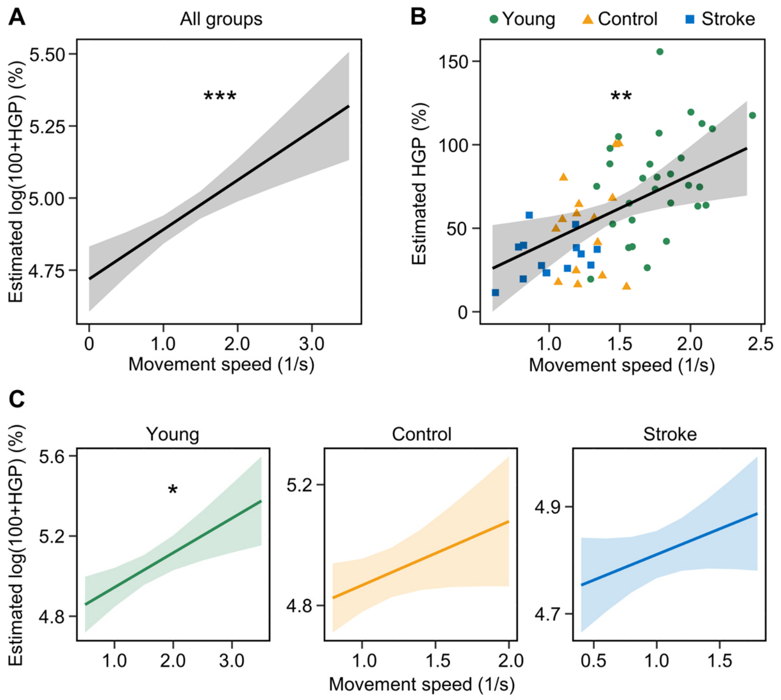


**Supplementary Figure 3 Relation between high-gamma power in PMC and movement speed.** (**A**) Effect plot of the fixed effect *movement speed* in a trial-level linear mixed-effects model across groups for the relation between *movement speed* and high-gamma power. (**B**) Effect plot of the fixed effect movement speed in a participant-level linear regression model across groups for the relation between *movement speed* and high-gamma power. (**C**) Effect plots of the fixed effect *movement speed* in trial-level mixed-effects models for the relation between *movement speed* and high-gamma power. Significance markers: *$p < 0.05$, **$p < 0.01$, ***$p < 0.001$

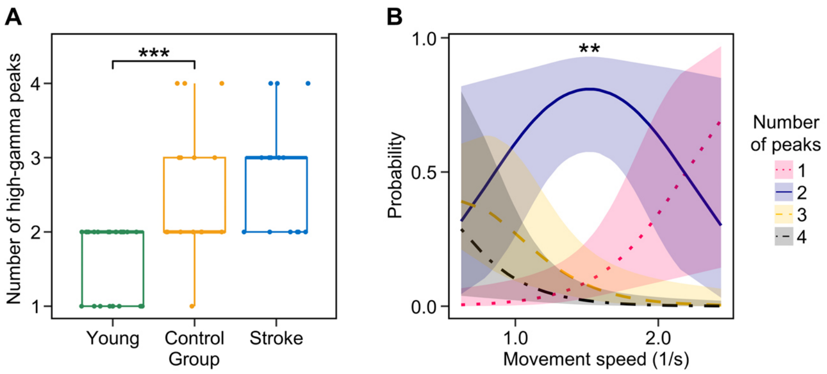


**Supplementary Figure 4 Number of high-gamma peaks in PMC.** (**A**) Distribution of the number of individual high-gamma peaks in PMC. The number of peaks was counted for participant mean of high-gamma power time courses. Asterisks indicate significant group differences. (**B**) Effect plot for the effect of the independent variable *movement speed* in a cumulative link model that models the relationship between the number of high-gamma peaks in PMC, *movement speed* and group with participant-level data from all three groups combined. The effect of *movement speed* was significant compared to a reduced model, the effect of group (not shown) was not significant. Significance markers: $^{**}p < 0.01$, $^{***}p < 0.001$

### Supplementary Tables

P-values reported in the Supplementary Tables are uncorrected. P-values after correction for multiple comparisons are reported in the results section of the main publication. Bold p-values reach statistical significance.

**Supplementary Table 1 Group differences in clinical scores and structural measures**

| Comparison | Variable | Statistical test | t | W | DF | p |
|---|---|---|---|---|---|---|
| Stroke – control | UEFM | WRST | | 30.5 | | **<0.001** |
| Stroke – control | ARAT | WRST | | 51.0 | | **0.004** |
| Stroke – control | BBT | t-test | -3.8 | | 19.2 | **0.001** |
| Stroke – control | NHPT | t-test | -2.7 | | 18.9 | **0.02** |
| Stroke – control | Grip force | WRST | | 64.0 | | 0.08 |
| Stroke – control | KG force | t-test | 0.4 | | 15.8 | 0.69 |
| Stroke – control | CST-FA | t-test | 1.13 | | 22.8 | 0.27 |

DF: degrees of freedom, WRST: Wilcoxon rank-sum test, KG force: key grip force

**Supplementary Table 2 Differences in movement speed and block-wise improvement**

| Comparison | Variable | Statistical test | t | DF | p |
|---|---|---|---|---|---|
| Stroke – control | Movement speed | t-test | -3.6 | 24.3 | **0.001** |
| Control – young | Movement speed | t-test | -7.4 | 40.8 | **<0.001** |
| Stroke B1 – stroke B6 | Movement speed | t-test | -2.4 | 24.8 | **0.02** |
| Control B1 – control B6 | Movement speed | t-test | -2.8 | 25.5 | **0.009** |
| Young B1 – young B6 | Movement speed | t-test | -3.8 | 55.6 | **<0.001** |
| Stroke – control | Block-wise improvement | t-test | 1.9 | 20.5 | 0.07 |
| Control - young | Block-wise improvement | t-test | -1.0 | 39.6 | 0.33 |

DF: degrees of freedom, B1: block 1, B6: block 6

**Supplementary Table 3 Group differences in high-gamma power**

| Comparison | Variable | Statistical test | t | DF | p |
|---|---|---|---|---|---|
| Stroke – control | HGP M1 | t-test | -2.2 | 18.9 | **0.04** |
| Stroke – control | HGP PMC | t-test | -2.3 | 19.4 | **0.03** |
| Control – young | HGP M1 | t-test | -2.3 | 34.4 | **0.03** |
| Control – young | HGP PMC | t-test | -2.9 | 30.1 | **0.008** |

DF: degrees of freedom, HGP: high-gamma power

**Supplementary Table 4 Linear regression models of high-gamma power, structure and clinical scores**

| | | | Model summary | | | | Whole model | | |
|---|---|---|---|---|---|---|---|---|---|
| **Group** | **DV** | **IVs** | **Estimate** | **SE** | **t** | **p** | **F** | **DF** | **p** |
| Stroke | HGP M1 | ChaCo M1 | -13.3 | 17.3 | -0.8 | 0.47 | 0.4 | 2, 7 | 0.71 |
| | | Performing hand (r) | 2.4 | 7.0 | 0.3 | 0.74 | | | |
| Stroke | HGP PMC | ChaCo PMC | -7.0 | 25.0 | -0.3 | 0.79 | 0.4 | 2, 7 | 0.71 |
| | | Performing hand (r) | 7.9 | 10.0 | 0.8 | 0.46 | | | |
| Stroke | HGP M1 | CST-FA | -0.3 | 0.6 | -0.5 | 0.64 | 0.8 | 2, 7 | 0.49 |
| | | Performing hand (r) | 0.07 | 0.06 | 1.2 | 0.25 | | | |
| Stroke | HGP PMC | CST-FA | -0.9 | 1.2 | -0.8 | 0.45 | 0.6 | 2, 7 | 0.58 |
| | | Performing hand (r) | 0.1 | 0.1 | 1.0 | 0.34 | | | |
| Stroke | HGP M1 | UEFM | -0.2 | 0.5 | -0.3 | 0.74 | 0.1 | 2, 11 | 0.94 |
| | | Performing hand (r) | 1.0 | 5.8 | 0.2 | 0.86 | | | |
| Stroke | HGP PMC | UEFM | 0.5 | 0.7 | 0.8 | 0.47 | 0.3 | 2, 11 | 0.71 |
| | | Performing hand (r) | 0.3 | 7.7 | 0.05 | 0.97 | | | |
| Stroke | HGP M1 | ARAT | -1.4 | 0.7 | -1.9 | 0.09 | 1.8 | 2, 11 | 0.21 |
| | | Performing hand (r) | 3.3 | 4.9 | 0.7 | 0.52 | | | |
| Stroke | HGP PMC | ARAT | -1.2 | 1.1 | -1.0 | 0.32 | 0.6 | 2, 11 | 0.56 |
| | | Performing hand (r) | 5.1 | 7.4 | 0.7 | 0.51 | | | |
| Stroke | HGP M1 | BBT | -0.06 | 0.2 | -0.3 | 0.80 | 0.04 | 2, 11 | 0.97 |
| | | Performing hand (r) | 1.5 | 7.0 | 0.2 | 0.84 | | | |
| Stroke | HGP PMC | BBT | 0.2 | 0.3 | 0.6 | 0.56 | 0.2 | 2, 11 | 0.78 |
| | | Performing hand (r) | -1.2 | 9.5 | -0.1 | 0.90 | | | |
| Stroke | HGP M1 | NHPT | -4.5 | 27.0 | -0.2 | 0.87 | 0.02 | 2, 11 | 0.99 |
| | | Performing hand (r) | 0.9 | 6.5 | 0.1 | 0.89 | | | |
| Stroke | HGP PMC | NHPT | 20.5 | 36.6 | 0.6 | 0.59 | 0.2 | 2, 11 | 0.81 |
| | | Performing hand (r) | -0.3 | 8.8 | -0.03 | 0.98 | | | |
| Stroke | HGP M1 | Grip force | -2.5 | 10.1 | -0.3 | 0.81 | 0.03 | 2, 11 | 0.97 |
| | | Performing hand (r) | 1.3 | 6.6 | 0.2 | 0.85 | | | |
| Stroke | HGP PMC | Grip force | -0.4 | 13.9 | -0.03 | 0.98 | 0.1 | 2, 11 | 0.94 |
| | | Performing hand (r) | 2.7 | 9.1 | 0.3 | 0.77 | | | |
| Stroke | HGP M1 | Key grip force | -2.9 | 12.2 | -0.2 | 0.82 | 0.03 | 2, 11 | 0.97 |
| | | Performing hand (r) | 0.9 | 5.9 | 0.2 | 0.88 | | | |
| Stroke | HGP PMC | Key grip force | -3.4 | 16.7 | -0.2 | 0.84 | 0.08 | 2, 11 | 0.92 |
| | | Performing hand (r) | 3.3 | 8.2 | 0.4 | 0.69 | | | |

DV: dependent variable, IVs: independent variables, SE: standard error, DF: degrees of freedom, HGP: high-gamma power. Performing hand indicates the effect of the right hand in comparison to the left hand.

**Supplementary Table 5 Linear mixed-effects models of high-gamma power and movement speed**

| | | | Model summary | | | Model comparison | | |
|---|---|---|---|---|---|---|---|---|
| **Group** | **DV** | **IVs** | **Estimate** | **SE** | **t** | **$X^2$** | **DF** | **p** |
| All groups | Log(100+HGP M1) | *Fixed effects* | | | | | | |
| | | Movement speed | 0.16 | 0.04 | 3.8 | 12.6 | 1 | **<0.001** |
| | | Group (stroke) | -0.04 | 0.05 | -0.9 | | | |
| | | Group (young) | 0.02 | 0.05 | 0.5 | | | |
| | | Performing hand (r) | -0.01 | 0.05 | -0.3 | | | |
| | | *Random effects* | **Variance** | **SD** | | | | |
| | | Participant (intercept) | 0.07 | 0.26 | | | | |
| | | Movement speed (Slope) | 0.05 | 0.23 | | | | |
| All groups | Log(100+HGP PMC) | *Fixed effects* | | | | | | |
| | | Movement speed | 0.17 | 0.04 | 4.1 | 14.2 | 1 | **<0.001** |
| | | Group (stroke) | -0.07 | 0.06 | -1.2 | | | |
| | | Group (young) | 0.05 | 0.06 | 0.9 | | | |
| | | Performing hand (r) | -0.05 | 0.06 | -0.8 | | | |
| | | *Random effects* | **Variance** | **SD** | | | | |
| | | Participant (intercept) | 0.09 | 0.30 | | | | |
| | | Movement speed (slope) | 0.05 | 0.22 | | | | |
| Stroke | Log(100+HGP M1) | *Fixed effects* | | | | | | |
| | | Movement speed | 0.09 | 0.06 | 1.5 | 2.2 | 1 | 0.14 |
| | | Performing hand (r) | -0.01 | 0.04 | -0.4 | | | |
| | | *Random effects* | **Variance** | **SD** | | | | |
| | | Participant (intercept) | 0.002 | 0.05 | | | | |
| Stroke | Log(100+HGP PMC) | *Fixed effects* | | | | | | |
| | | Movement speed | 0.10 | 0.06 | 1.5 | 2.3 | 1 | 0.13 |
| | | Performing hand (r) | -0.003 | 0.05 | -0.1 | | | |
| | | *Random effects* | **Variance** | **SD** | | | | |
| | | Participant (intercept) | 0.01 | 0.08 | | | | |
| Control | Log(100+HGP M1) | *Fixed effects* | | | | | | |
| | | Movement speed | 0.30 | 0.13 | 2.7 | 4.3 | 1 | **0.04** |
| | | Performing hand (r) | 0.02 | 0.07 | 0.2 | | | |
| | | *Random effects* | **Variance** | **SD** | | | | |
| | | Participant (intercept) | 0.18 | 0.43 | | | | |
| | | Movement speed (slope) | 0.17 | 0.41 | | | | |
| Control | Log(100+HGP PMC) | *Fixed effects* | | | | | | |
| | | Movement speed | 0.21 | 0.12 | 1.8 | 2.8 | 1 | 0.10 |
| | | Performing hand (r) | -0.08 | 0.09 | -0.9 | | | |
| | | *Random effects* | **Variance** | **SD** | | | | |
| | | Participant (intercept) | 0.17 | 0.41 | | | | |
| | | Movement speed (slope) | 0.12 | 0.35 | | | | |
| Young | Log(100+HGP M1) | *Fixed effects* | | | | | | |
| | | Movement speed | 0.13 | 0.05 | 2.4 | 5.1 | 1 | **0.02** |
| | | *Random effects* | **Variance** | **SD** | | | | |
| | | Participant (intercept) | 0.11 | 0.33 | | | | |
| | | Movement speed (slope) | 0.05 | 0.22 | | | | |
| Young | Log(100+HGP PMC) | *Fixed effects* | | | | | | |
| | | Movement speed | 0.17 | 0.06 | 3.1 | 8.0 | 1 | **0.005** |
| | | *Random effects* | **Variance** | **SD** | | | | |
| | | Participant (intercept) | 0.15 | 0.40 | | | | |
| | | Movement speed (slope) | 0.05 | 0.23 | | | | |

DV: dependent variable, IVs: independent variables, SE: standard error, SD: standard deviation, DF: degrees of freedom, HGP: high-gamma power. Performing hand indicates the effect of the right hand in comparison to the left hand, Group indicates the effect of the respective group in comparison to the control group. Model comparison against a reduced model without the fixed effect in question.

**Supplementary Table 6 Linear regression models of high-gamma power and movement speed**

| | | | Model summary | | | | Whole model | | |
|---|---|---|---|---|---|---|---|---|---|
| **Group** | **DV** | **IVs** | **Estimate** | **SE** | **t** | **p** | **F** | **DF** | **p** |
| All groups | HGP M1 | Movement speed | 34.7 | 12.4 | 2.8 | **0.007** | 7.1 | 4, 53 | **<0.001** |
| | | Group (stroke) | -4.3 | 8.8 | -0.5 | 0.63 | | | |
| | | Group (young) | 4.0 | 10.5 | 0.4 | 0.71 | | | |
| | | Performing hand (r) | -5.6 | 8.5 | -0.7 | 0.51 | | | |
| All groups | HGP PMC | Movement speed | 40.0 | 14.5 | 2.7 | **0.008** | 10.2 | 4, 53 | **<0.001** |
| | | Group (stroke) | -7.0 | 10.3 | -0.7 | 0.50 | | | |
| | | Group (young) | 15.0 | 12.2 | 1.2 | 0.23 | | | |
| | | Performing hand (r) | -11.9 | 9.9 | -1.2 | 0.24 | | | |

DV: dependent variable, IVs: independent variables, SE: standard error, DF: degrees of freedom, HGP: high-gamma power. Performing hand indicates the effect of the right hand in comparison to the left hand, Group indicates the effect of the respective group in comparison to the control group.

**Supplementary Table 7 Group differences in high-gamma peak frequency**

| Comparison | Variable | Statistical test | W | p |
|---|---|---|---|---|
| Stroke – control | HGF M1 | WRST | 107.0 | 0.95 |
| Stroke – control | HGF PMC | WRST | 100.5 | 0.86 |
| Control – young | HGF M1 | WRST | 46.5 | **<0.001** |
| Control – young | HGF PMC | WRST | 55.0 | **<0.001** |

HGF: high-gamma peak frequency, WRST: Wilcoxon rank-sum test

**Supplementary Table 8 Linear regression models of high-gamma peak frequency and movement speed**

| | | | Model summary | | | | Whole model | | |
|---|---|---|---|---|---|---|---|---|---|
| **Group** | **DV** | **IVs** | **Estimate** | **SE** | **t** | **p** | **F** | **DF** | **p** |
| All groups | HGF M1 | Movement speed | 5.0 | 3.4 | 1.5 | 0.15 | 7.3 | 4, 53 | **<0.001** |
| | | Group (stroke) | 2.6 | 2.4 | 1.1 | 0.28 | | | |
| | | Group (young) | 7.1 | 2.9 | 2.5 | **0.02** | | | |
| | | Performing hand (r) | -1.4 | 2.3 | -0.6 | 0.56 | | | |
| All groups | HGF PMC | Movement speed | 5.8 | 3.1 | 1.9 | 0.07 | 7.0 | 4, 53 | **<0.001** |
| | | Group (stroke) | 2.1 | 2.2 | 1.0 | 0.33 | | | |
| | | Group (young) | 4.5 | 2.6 | 1.7 | 0.09 | | | |
| | | Performing hand (r) | -0.1 | 2.1 | -0.05 | 0.96 | | | |

DV: dependent variable, IVs: independent variables, SE: standard error, DF: degrees of freedom, HGF: high-gamma peak frequency. Performing hand indicates the effect of the right hand in comparison to the left hand, Group indicates the effect of the respective group in comparison to the control group.

**Supplementary Table 9 Group differences in the number of high-gamma peaks**

| Comparison | Variable | Statistical test | W | p |
|---|---|---|---|---|
| Stroke – control | nPeaks M1 | WRST | 127.5 | 0.30 |
| Stroke – control | nPeaks PMC | WRST | 117.5 | 0.58 |
| Control – young | nPeaks M1 | WRST | 314.0 | **0.009** |
| Control – young | nPeaks PMC | WRST | 352.5 | **<0.001** |

nPeaks: number of high-gamma peaks, WRST: Wilcoxon rank sum test

**Supplementary Table 10 Cumulative link models of the number of high-gamma peaks, movement speed and group**

| | | | Model summary | | | Model comparison | | |
|---|---|---|---|---|---|---|---|---|
| **Group** | **DV** | **IVs** | **Estimate** | **SE** | **z** | **$X^2$** | **DF** | **p** |
| All groups | nPeaks M1 | Movement speed | -4.37 | 1.38 | -3.2 | 12.1 | 1 | **<0.001** |
| | | Group | | | | 0.03 | 2 | 0.98 |
| | | Group (stroke) | -0.13 | 0.78 | -0.2 | | | |
| | | Group (young) | -0.10 | 0.99 | -0.1 | | | |
| | | Performing hand (r) | 0.15 | 0.76 | 0.2 | | | |
| All groups | nPeaks PMC | Movement speed | -3.33 | 1.31 | -2.5 | 7.4 | 1 | **0.006** |
| | | Group | | | | 2.5 | 2 | 0.29 |
| | | Group (stroke) | -0.33 | 0.80 | -0.4 | | | |
| | | Group (young) | -1.87 | 1.30 | -1.4 | | | |
| | | Performing hand (r) | -0.75 | 0.77 | -1.0 | | | |

DV: dependent variable, IVs: independent variables, SE: standard error, DF: degrees of freedom, nPeaks: number of high-gamma peaks. Performing hand indicates the effect of the right hand in comparison to the left hand, Group indicates the effect of the respective group in comparison to the control group. Model comparison against a reduced model without the fixed effect in question.

**Supplementary Table 11 Linear mixed-effects models of high-gamma power and trial-wise improvement**

| | | | Model summary | | | Model comparison | | |
|---|---|---|---|---|---|---|---|---|
| **Group** | **DV** | **IVs** | **Estimate** | **SE** | **t** | **$X^2$** | **DF** | **p** |
| All groups | Log(100+HGP M1) | *Fixed effects* | | | | | | |
| | | Trial-wise improvement | -0.0002 | 0.0006 | -0.3 | 0.08 | 1 | 0.77 |
| | | Movement speed | 0.15 | 0.04 | 3.4 | | | |
| | | Group (stroke) | -0.04 | 0.05 | -0.9 | | | |
| | | Group (young) | 0.03 | 0.05 | 0.6 | | | |
| | | Performing hand (r) | -0.006 | 0.05 | -0.1 | | | |
| | | *Random effects* | **Variance** | **SD** | | | | |
| | | Participant (intercept) | 0.06 | 0.24 | | | | |
| | | Movement speed (slope) | 0.04 | 0.21 | | | | |
| All groups | Log(100+HGP PMC) | *Fixed effects* | | | | | | |
| | | Trial-wise improvement | 0.0004 | 0.0006 | 0.7 | 0.5 | 1 | 0.48 |
| | | Movement speed | 0.20 | 0.03 | 6.5 | | | |
| | | Group (stroke) | -0.06 | 0.06 | -1.0 | | | |
| | | Group (young) | 0.03 | 0.06 | 0.6 | | | |
| | | Performing hand (r) | -0.06 | 0.06 | -0.9 | | | |
| | | *Random effects* | **Variance** | **SD** | | | | |
| | | Participant (intercept) | 0.03 | 0.16 | | | | |

DV: dependent variable, IVs: independent variables, SE: standard error, SD: standard deviation, DF: degrees of freedom, HGP: high-gamma power. Performing hand indicates the effect of the right hand in comparison to the left hand, Group indicates the effect of the respective group in comparison to the control group. Model comparison against a reduced model without the fixed effect in question.

**Supplementary Table 12 Linear regression models of high-gamma power and block-wise improvement**

| | | | Model summary | | | | Whole model | | |
|---|---|---|---|---|---|---|---|---|---|
| **Group** | **DV** | **IVs** | **Estimate** | **SE** | **t** | **p** | **F** | **DF** | **p** |
| All groups | HGP M1 | Block-wise improvement | -0.4 | 0.3 | -1.5 | 0.13 | 8.5 | 5, 52 | **<0.001** |
| | | Movement speed block 1 | 43.0 | 13.1 | 3.3 | **0.002** | | | |
| | | Group (stroke) | 1.3 | 8.3 | 0.2 | 0.88 | | | |
| | | Group (young) | 4.5 | 9.8 | 0.5 | 0.65 | | | |
| | | Performing hand (r) | -7.1 | 7.9 | -0.9 | 0.37 | | | |
| All groups | HGP PMC | Block-wise improvement | -0.3 | 0.3 | -0.9 | 0.38 | 9.7 | 5, 52 | **<0.001** |
| | | Movement speed block 1 | 47.5 | 16.0 | 3.0 | **0.005** | | | |
| | | Group (stroke) | -2.6 | 10.1 | -0.3 | 0.80 | | | |
| | | Group (young) | 15.7 | 12.0 | 1.3 | 0.20 | | | |
| | | Performing hand (r) | -13.1 | 9.6 | -1.4 | 0.18 | | | |

DV: dependent variable, IVs: independent variables, SE: standard error, DF: degrees of freedom, HGP: high-gamma power. Performing hand indicates the effect of the right hand in comparison to the left hand, Group indicates the effect of the respective group in comparison to the control group.

### Supplementary Methods

#### Fractional Anisotropy of the corticospinal tract

*Preprocessing*

Anatomical data preprocessing

The T1-weighted (T1w) image was corrected for intensity non-uniformity using N4BiasFieldCorrection[1] (ANTs 2.4.0), and used as T1w-reference throughout the workflow. The T1w-reference was then skull-stripped using antsBrainExtraction.sh (ANTs 2.4.0), using OASIS as target template. Spatial normalization to the ICBM 152 Nonlinear Asymmetrical template version 2009c[2] (RRID:SCR_008796) was performed through nonlinear registration with antsRegistration[3] (ANTs 2.4.0, RRID:SCR_004757), using brain-extracted versions of both T1w volume and template. Brain tissue segmentation of cerebrospinal fluid, white-matter and grey-matter was performed on the brain-extracted T1w using FAST[4] (FSL 6.0.5.1:57b01774, RRID:SCR_002823).

Diffusion data preprocessing

Any images with a b-value less than 100 s/mm$^2$ were treated as a b=0 image. Marchenko-Pastur Principal Component Analysis (MP-PCA) denoising as implemented in MRtrix3's dwidenoise[5] was applied with a 5-voxel window. After MP-PCA, B1 field inhomogeneity was corrected using dwibiascorrect from MRtrix3 with the N4 algorithm.[1] After B1 bias correction, the mean intensity of the diffusion-weighted imaging (DWI) series was adjusted so all the mean intensity of the b=0 images matched across each separate DWI scanning sequence.

FSL (version 6.0.5.1:57b01774)'s eddy was used for head motion correction and Eddy current correction.[6] Eddy was configured with a q-space smoothing factor of 10, a total of 5 iterations, and 1000 voxels used to estimate hyperparameters. A linear first level model and a linear second level model were used to characterize Eddy current-related spatial distortion. q-space coordinates were forcefully assigned to shells. Field offset was attempted to be separated from subject movement. Shells were aligned post-eddy. Eddy's outlier replacement was run.[7] Data were grouped by slice, only including values from slices determined to contain at least 250 intracerebral voxels. Groups deviating by more than 4 standard deviations from the prediction had their data replaced with imputed values. Data was collected with reversed phase-encode blips, resulting in pairs of images with distortions going in opposite directions. Here, b=0 reference images with reversed phase encoding directions were used along with an equal number of b=0 images extracted from the DWI scans. From these pairs the susceptibility-induced off-resonance field was estimated using a method similar to that described in Andersson et al.[8] The fieldmaps were ultimately incorporated into the Eddy current and head motion correction interpolation. Final interpolation was performed using the jac method.

Several confounding time-series were calculated based on the preprocessed DWI: framewise displacement using the implementation in Nipype (following the definitions by Power et al.[9]). The head-motion estimates calculated in the correction step were also placed within the corresponding confounds file. Slicewise cross correlation was also calculated. The DWI time-series were resampled to anterior commissure - posterior commissure (ACPC), generating a preprocessed DWI run in ACPC space with 2 mm isotropic voxels.

Many internal operations of QSIPrep use Nilearn 0.9.2[10] (RRID:SCR_001362) and Dipy.[11] For more details of the pipeline, see the section corresponding to workflows in QSIPrep's documentation.
